# Pooled genetic screens identify breast cancer risk genes involved in evasion from T cell-mediated killing

**DOI:** 10.1101/2024.05.10.593465

**Authors:** Wei Shi, Yi Luo, Jacqueline M. Burrows, Debra Black, Andrew Civitarese, Laura Perlaza-Jimenez, Ping Zhang, Murray Manning, Natasha Tuano, Miguel E. Rentería, Christos Xiao, Siok-Keen Tey, Joseph Rosenbluh, Corey Smith, Georgia Chenevix-Trench, Jonathan Beesley

**Affiliations:** Cancer Division, QIMR Berghofer, Brisbane, QLD 4006, Australia; Infection and Immunology Division, QIMR Berghofer, Brisbane, QLD 4006, Australia; Cancer Research Program and Department of Biochemistry and Molecular Biology, Biomedicine Discovery Institute, Monash University, Clayton, VIC, Australia; Monash Functional Genomics Platform, Monash University, Clayton, VIC 3800, Australia; Mental Health & Neuroscience Program, QIMR Berghofer Medical Research Institute, Brisbane QLD, 4006, Australia

## Abstract

Genome-wide association studies have identified more than 220 loci associated with breast cancer susceptibility. A major challenge is now to identify the effector genes with plausible functions in the context of breast cancer risk. We have previously performed pooled CRISPR screens to identify target genes at risk loci that drive cancer hallmarks including proliferation or modulating DNA damage response. We now extend these screens to identify genes involved in response to cytotoxic T lymphocyte (CTL) killing. We performed knockout and inhibition screens to identify genes that affect the response of the MCF7 human breast cancer cell line to CTL killing in an *in vitro* co-culture system. We identified 33 candidate risk genes associated with resistance or sensitisation to T cell-mediated killing. Using single gene perturbation, we showed that deletion of candidate risk genes *IRF1, ATF7IP, CCDC170* and *CASP8* induced resistance, while ablation of *CFLAR, CREBBP*, and *PRMT7* sensitized cells to CTL killing. We used reporter assays to show that the risk-associated alleles at rs736801 and rs3769821 reduced transactivation of the *IRF1* and *CASP8* promoters, respectively. We showed that pharmacological inhibition of PRMT7 rendered breast cells sensitive to CTL killing and *PRMT7* levels were negatively correlated with CD8+ infiltration and patient survival in luminal A breast cancer patient cohorts. Our results demonstrate that phenotypic pooled CRISPR screens are a useful approach for high throughput functional follow-up of GWAS findings, identifying genes which alter immune responses to breast cancer which offer opportunities to enhance immunotherapy.

## Introduction

Genome-wide association studies (GWAS) have identified over 220 loci associated with breast cancer risk ^1,2^. However, the pathway to translate genetic discoveries faces major challenges. Genetic fine-mapping and computational analyses have predicted more than 1200 target genes, including 191 with high confidence ^3^. We previously showed that high-throughput custom CRISPR screens are effective at identifying breast cancer risk genes involved in cellular proliferation, tumorigenicity and DNA damage repair ^4^. In addition to these cancer hallmarks, pathway analysis of the predicted target genes showed that immune system processes were among the top enriched pathways for both estrogen receptor (ER)-positive and ER-negative GWAS ^3^, warranting a screen to determine whether any of these genes is involved in immunosurveillance.

Cancer immunoediting consists of three phases: elimination (also known as cancer immunosurveillance), equilibrium, and escape, during which the immune system not only protects against cancer development but also shapes the character of developing tumors ^5–9^. Cancer immunosurveillance recognizes neo-antigens on transformed cells and eliminates them before tumors are detectable. This process, involving components of the innate and adaptive immune systems, is essential in early anti-tumor responses to induce cell death and increase tumor immunogenicity ^5,6,8^. Cytotoxic T cells are capable of antigen-directed target cell killing and are key drivers of cancer immunoediting ^10^. While much focus has been placed on understanding the immune escape phase in the context of breast cancer, it is possible that the poorly understood elimination phase is likely to involve cancer risk genes which act in earlier phases of tumorigenesis ^11^. Immune surveillance depends on several factors including cytokine production (e.g. IFNγ and TNFα), and effector molecules (e.g. granzymes and perforin) on immune cells, to effectively eliminate cancer cells.

There have been few studies to date on immune functions in women genetically predisposed to breast cancer. A recent single cell transcriptomic analysis of risk reduction mammoplasties showed that immune cells from *BRCA1/2* carriers had a distinct gene expression signature indicative of potential immune exhaustion, suggesting that immune escape mechanisms could manifest in non-cancerous tissues during very early stages of tumor initiation ^12,13^. In addition, a recent study showed that the anti-tumor response in women becomes less active during the transition from ductal carcinoma in situ (DCIS) to invasive ductal breast cancer (IDC), with fewer activated granzyme B-expressing CD8+ T cells in IDC than DCIS ^14^. They reasoned that in DCIS, with the help of a mostly intact physical barrier, the immune cells can easily eliminate the relatively few cancer cells that are exposed and keep the tumor in check. Therefore, the elimination process is critical to foil the DCIS-IDC transition, with clinically detected IDC representing a failure of elimination. Furthermore, several strains of immunodeficient mice develop spontaneous mammary carcinomas, demonstrating the importance of elimination ^5,15^.

We hypothesized that some genes implicated by GWAS of breast cancer risk are involved in modulating the sensitivity of breast cancer cells to immune killing. We therefore performed pooled CRISPR screens in a HER2-expressing human breast cancer cell line, co-cultured with HLA-matched, HER2-restricted T cell receptor (TCR)-expressing T cells. These screens identified novel resistor and sensitizer roles for predicted breast cancer risk genes, including genes directly regulated by breast cancer risk variants, and the pharmacologically modulated target PRMT7.

## Material and methods

### Identification of appropriate breast cell lines for the CTL killing assay

To establish the CTL killing assay with breast cell lines, we first evaluated HER2 (*ERBB2*) expression in the breast cell lines using RNA-seq data from Cancer Cell Line Encyclopedia ^16^ and from published data for B80-T5 ^4^, and validated by flow cytometry. We determined the HLA-type of the HER2+ cell lines and found an HLA-matching donor as a source of T cells. Breast cell line DNAs were genotyped at the Genomic Core Facility of Erasmus University Rotterdam (Netherlands) using the Illumina Global Screening Array. We used the SNP2HLA method ^17^ to impute classical HLA types based on the observed genotypes from the SNP array.

### Isolation of peripheral blood mononuclear cells

Blood samples from healthy donors were collected with donors’ informed consent according to the requirements of the Human Research Ethics Committee of QIMR Berghofer. The donors were HLA-typed (Victorian Transplantation & Immunogenetics Services, Australian Red Cross Lifeblood, Melbourne, Australia) to identify donors, and peripheral blood mononuclear cells (PBMCs) from a female healthy HLA-A2 donor (to match the breast cell lines we used for the screen) were isolated using Ficoll separation. Whole blood from EDTA tubes (BD 366643) was centrifuged at 500g for 10 minutes at room temperature with no brake to remove plasma, the remaining blood was mixed at a 1:1 ratio with room temperature RPMI medium, layered over Ficoll-Paque PLUS (GE 17144002) or Lymphoprep™ (stemcell Cat#07851) in SepMate tube (Stemcell Cat#85450), and spun at 1200g for 10 minutes at room temperature. The PBMC layer was collected and washed with RPMI. Cells were counted, centrifuged again and resuspended in freezing medium (90% fetal bovine serum (FBS) + 10% DMSO) and stored in cryogenic vials in liquid nitrogen.

### Generation of HER2-T cell receptor expressing T cells

To establish an antigen-specific CTL killing assay of breast cell lines, we firstly generated HER2-restricted T cell receptor (TCR)-expressing T cells. We used a lentivirus-based allo-restricted TCR-expressing construct for the HER2-derived peptide 369 (HER_369_)^18^. The TCR was modified within the constant regions to enhance stability and expression levels by exchange of selected amino acids with the murine counterparts. ^19^. The TCR α- and β-chain sequences were then synthetically introduced into a lentivirus plasmid as one transcript using a cleavage protein to produce the α- and β-chains as two products in equal ratios driven by the promoter human elongation factor-1 alpha (EF-1 alpha) (Biosettia Inc., San Diego). This construct (pLV-EF1a-HER2-1-TCR1), along with plasmids pMDL, pVSV-G and pREV, were transfected into HEK293T packaging cells. Lentiviral supernatant was harvested after 24, 48 and 72 hour incubation in DMEM containing 10% FBS and passed through a 0.45-μm Milli-hex filter before untra-centrifugation (10500 rpm, > 6 hr at 4 degrees).

To activate the T cells in PBMCs, 24-well plates (Falcon, Franklin Lakes, NJ) were coated with a mixture of anti-Human-CD3 and CD28 (clone OKT3 and CD28.2, eBioscience) mAbs at 0.5 µg/ml. PBMCs were plated at 2x10^6^ per well to the CD3/CD8 coated plate in AIMV complete media (AIM V™ SFM, Gibco Cat No: 0870112DK + 5% CTS™ Immune Cell SR, Gibco Cat No: A2596101) supplemented with 300 IU/ml rIL-2. Eighteen hours after activation, PBMCs were transduced with the HER2-TCR expressing lentivirus using the spin infection method on CD3/CD28 and Retronectin (16 μg/ml) coated plates with the addition of 4 μM TBK1/IKKɛ complex inhibitor BX-795 to enhance lentiviral delivery ^20^. Six hours after infection, the media was adjusted to 1.3 μM BX-795. After a further 60 hours, the media was changed to RPMI with 10% heat inactivated FBS and the cells plated into G-Rex^®^ 6 Well plates (Wilson Wolf P/N 80240M) as per manufacturer’s instructions for large scale expansion before harvesting the cells on day 13. A single batch of HER2-TCR T cells (henceforth called HER2-restricted T cells) was generated, expanded and cryo-preserved for all screens and subsequent validations. The HER2-TCR infection rate was determined by flow cytometry in CD3+, CD4+ and CD8+ cells with a TCR Vβ8 antibody, in comparison with mock-transduced PBMCs. The panel included antibodies for TCR Vβ8 FITC (clone 56C5.2, Beckman Cat# IM1233), CD3 BV711 (clone SK7, BD Pharmingen™ Cat# 740832), CD4 (clone SK3, BD Pharmingen™ Cat# 612748), CD8 (clone SK1, BD Pharmingen™ Cat# 565310) and viability dye Sytox Blue (Invitrogen™ S34857) and was analysed on BD LSRFortessa™ Cell Analyzer.

### T cell cytotoxicity assay

In order to perform the pooled CRISPR screens in breast cells, we adopted the two cell type assay previously described to identify essential genes for immunotherapy ^21–23^. HLA-A2-positive breast carcinoma cell lines, MCF-7, SKBR3, MDA-MB-231 and the immortalised normal breast cell line B80-T5 ^24^ were cultured in 96 well E-Plates (ACEA Biosciences) for 24 hours followed by co-culturing with HLA-A2-HER2-restricted T cells on the xCELLigence RTCA instrument (ACEA Biosciences) until the cytolysis plateaus ^25,26^. We added 100 uL of RPMI1640 containing 10% FBS to each well, and measured the background impedance, displayed as the Cell Index. Dissociated breast cells were then seeded at a density of 20,000 (MCF7, SKBR3 and MDA-MD-231) or 10,000 (B80-T5) cells/well of the E-Plate in a volume of 100 uL RPMI-10% FBS media and allowed to passively adhere on the electrode surface for 30 minutes inside the cell culture incubator hosting the RTCA instrument. Impedance data, Cell Index, Normalised Cell Index and percentage cytolysis were recorded at 15 minute intervals and displaced in real-time for the entire duration of the experiment. The initial cell index for each well before co-culture was the readout of breast cell health status, viability and growth. When the co-culture started, HER2-restricted T (effector) cells were added to the breast (target) cells at the indicated (E:T) ratio. Percent cytolysis was determined at each time point using the xIMT software (ACEA Biosciences). The rate of cytolysis was calculated for each well at every time point, using the Normalised Sample Cell Index and the Normalised Average Target Alone Control according to the following equation:

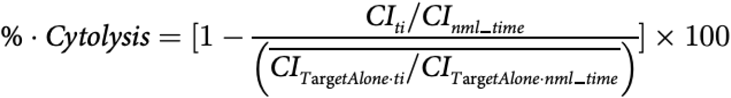

Effector cell only controls, full lysis controls and target plus mock-transduced PBMCs (T blast) controls were also set up during the optimization of the assay. Effector cell only controls were comparable to baseline level of impedance while full lysis controls show 100% cytolysis shortly after being applied to the adherent breast cells.

We assessed HER2 expression level in MCF7, SKBR3, MDA-MB-231 and B80-T5 by flow cytometry (Biolegend, PE anti-human CD340 (erbB2/HER-2) Antibody, Cat# 324406). We titrated the concentration of IFNγ ranging from 0.1 to 100 ng/ml to find the optimal low dose for enhancing MHC-I expression in MCF7 cells within 24 hr. The HLA-ABC level of MCF7 cells was assessed by flow cytometry (BD Pharmingen™ PE Mouse Anti-Human HLA-ABC, Cat# 565291) following 24hr treatment with different concentrations of IFNγ.

All cell lines were maintained in a humidified incubator at 37 °C with 5% CO2, and tested regularly for mycoplasma infection. Cell line identity for established cells was confirmed by Short Tandem Repeat analysis at the QIMR Berghofer Analytical Facility.

### Generation of lentiviruses

Lentiviruses were produced as previously described ^4^. Briefly, sgRNAs or expression vectors were transfected along with pMD2.G (Addgene #12259) and psPAX2 (Addgene #12260) lentiviral vectors which were into HEK293T packaging cells. Lentiviral supernatant was harvested after 48-h incubation in DMEM containing 30% FBS and passed through a 0.45-μm Milli-hex filter.

### Pooled sgRNA libraries

The custom CRISPRko (knockout) and CRISPRi (inhibition) screen libraries of candidate breast cancer risk genes were generated as described ^4^. Pooled oligonucleotides were cloned into the BsmBI sites of pLeniGuide-Puro (Addgene #52963). We included as positive controls core cell essential genes and genes known to confer resistance to T cell-mediated killing, including *APLNR, ARID2, B2M, BBS1, CD58, CTLA4, ERAP1, ERAP2, HLA-A, IFNGR1, IFNGR2, IRF2, IRF8, JAK1, JAK2, NLRC5, PBRM1, PDCD1, CD274, STAT1, TAP1, TAP2* and *TPP2* ^21,23,27,28^. All sgRNA sequences are available in Table S2. For each gene, we chose top scoring five sgRNAs designed using the CRISPick algorithm. Libraries were electroporated into NEB5α electrocompetent cells (NEB), grown overnight and plasmid DNA extracted using Qiagen Maxi Prep. For each library preparation, at least 1000X guide representation was ensured by counting surviving colonies from the serially diluted samples.

### Pooled CRISPR screens for resistance to T cell cytotoxicity

For CRISPR screens and validations, breast cell lines were transduced with Lenti-Cas9-2A-Blast (Addgene #73310) or Lenti-dCas9-KRAB-Blast (Addgene #89567). MCF7 cells stably expressing Cas9 or KRAB-dCas9 were established by transduction of the virus at MOI<0.7 ^29^, selected in 10 μg/ml blasticidin and maintained in 5 μg/ml blasticidin. MCF7 cells were transduced with CRISPRko or CRISPRi library virus at a MOI of 0.3 to ensure one guide per cell and obtain 1000 cells/sgRNA. Library infection was carried out in triplicate. Twenty-four hours after infection, cells were selected using puromycin (1 μg/ml) for 7 days. In order to assay for resistance to HER2-T cell cytotoxicity, library-infected MCF7 cells were cultured for 24 hr with interferon-gamma (IFNγ) at 10 ng/ml followed by co-culturing with HER2-restricted T cells at an Effector (E; HER2-T cells): Target (T; MCF7) ratio of 1:1. The control group was cultured for a recovery period equal to the T cell co-culture period. A parallel xCELLigence assay was run for each screen for real time monitoring of cytolysis. When the cytolysis rate reached 70% in the co-culture flask (MCF7+T cells), the media was removed from both the co-culture and the unexposed controls. After three washes with PBS, the cells were maintained with blasticidin (10 ug/ml) and puromycin (1 ug/ml) for another 48 hours. The cells were then harvested and genomic DNA was extracted using NucleoSpin Blood XL kit (Clontech). Libraries were prepared and sequenced as previously described ^4^.

### Pooled screen analysis

Raw sequencing reads were processed by trimming adaptors and vector sequences using *cutadapt* v1.13 with parameter ‘*-g TTGTGGAAAGGACGAAACACCG*’. sgRNA barcodes were counted using *MAGeCK* (v0.5.9.4) and the significance of treatment-dependent abundances calculated using maximum-likelihood estimation implemented in the MLE module, with 10 rounds of permutation (each round corresponding to 2 × gene number), correcting for MCF7 gene copy number using CNV array data from the Broad Institute Cancer Cell Line Encyclopedia and normalizing by counts of negative control sgRNAs. Experimental treatment conditions for MLE modelling were supplied as binary design matrices including variables for baseline, proliferation (to capture general fitness effects in culture), IFNγ response, and T cell exposure.

### Generation of stable cell lines for single gene validation

For single gene validation, the two top scoring sgRNAs were cloned into BsmBI-digested lentiGuide-Puro vector (Addgene #52963) (Table S3). MCF7 or B80-T5 cells with single gene perturbation were generated as previously described ^4^. Gene perturbation was confirmed by Western blotting or RT-qPCR. Total protein or RNA were isolated from these cells at the comparable passage number with which the cytolysis assay was performed. Total RNA isolation was performed using the RNeasy Mini Kit (Qiagen). cDNA was synthesised using the Maxima H Minus First Strand cDNA Synthesis Kit (Thermo Scientific) and amplified using PowerUp™ SYBR™ Green Master Mix (Thermo Scientific). The mRNA levels for each sample was measured in technical triplicates for each primer set, along with three housekeeping genes (*ACTIN, GUSB*, and *PUM1*; Table S4). Experiments were performed using an ABI ViiA(TM) 7 or ABI Q5 Quantstudio System (Applied Biosystems), and data processed using ABI QuantStudio™ Software (Applied Biosystems). The average Cт of target genes was compared with the geometric mean of three housekeeping genes to calculate ΔCт. Relative quantitation of each target gene was normalised to the corresponding sgRFP control line using the comparative Cт method (ΔΔCт).

Whole cell lysates were prepared with RIPA buffer containing protease inhibitors (Invitrogen) as per manufacturer’s instructions before quantification with BCA assay (Thermo Scientific). Samples were reduced and boiled prior to loading 20 μg per lane onto precast 4-15% Mini-Protean TGX polyacrylamide gels (BioRad Laboratories Inc) followed by transfer onto low-fluorescence PVDF membranes with a Trans-blot Turbo Transfer System (BioRad Laboratories Inc). After blocking for 1hr in Intercept (TBS) Blocking Buffer (LI-COR Biosciences), membranes were probed overnight at 4°C with primary antibodies against the respective target: ATF7IP (HPA023505, Sigma-Aldrich), CFLAR (#56343, Cell Signaling Technology), CREBBP CBP(D6C5) (#7389, Cell Signaling Technology), IFNGR2 (OTI1C2) (TA506734S, Origene Technologies Inc), IRF1 (VPA00801, BioRad Laboratories Inc) and PRMT7(D1K6R) (#14762, Cell Signaling Technology). Fluorescent detection and imaging on the LI-COR Odyssey CLx instrument (LI-COR Biosciences) was carried out the next day. All targets were normalized to a housekeeping gene, either beta-Actin (AC-74) (A2228, Sigma-Aldrich) or Cytochrome C oxidase (COXIV; 926-42214, LI-COR Biosciences), using the ImageStudio software package.

### Reporter constructs and assays

The *IRF1* (hg38 region chr5:132490551-132491102) and *CASP8* promoter containing rs3769821 (hg38 region chr2:201257548-201259062) were PCR amplified and cloned using Gibson assembly into the HindIII-linearised pGL3-Basic luciferase reporter (Table S5). The putative regulatory region (PRE) containing candidate causal risk variants (CCVs) upstream of *IRF1* (rs736801, hg38 region chr5:132497660-132498158) were identified based on ENCODE ChIP-seq for the H3K27ac active regulatory histone mark. The fragment was amplified using PCR primers listed in Table S5 and cloned using Gibson assembly into the Sall site of the pGL3-IRF1 promoter construct. Heterozygous genomic DNA was used to generate fragments containing reference or alternate alleles. All constructs were verified using Sanger sequencing. We transiently co-transfected breast cells with pGL3 luciferase reporter constructs and Renilla internal control plasmid. After 24 hours, cell lysates were prepared using the Dual-Glo Luciferase Kit (Promega) and luciferase luminescent activity measured with a Synergy H4 plate reader (Biotek). Luciferase activity was normalized to Renilla to correct for differences in transfection efficiencies or lysate preparation. Allele-specific activities were analysed after log transformation of normalised readings with two-way ANOVA followed by Dunnett’s test for multiple comparisons.

### Survival analysis

Bulk RNA-seq, molecular sub-type and survival data were obtained from METABRIC ^30^ and TCGA ^31^. Patients with multiple tumour samples were removed. The prognostic association with *PRMT7* gene expression was tested using Kaplan-Meier plots. To further test the survival association with *PRMT7* gene expression on the infiltration of CD8 positive T cells, quanTIseq ^32^ was used to quantify the fractions of immune cells in TCGA, the cohort was divided into four sub-groups (high or low *PRMT7* expression combined with high or low CD8 positive T cells infiltration) accordingly for Kaplan-Meier plots. The significance of survival difference was estimated using a log-rank test. All survival analyses including determining optimal cut-off points were performed using the R packages “survival” and “survminer”.

### In vitro PRMT7 inhibitor studies

Background impedance was measured by adding 100 µL of RPMI1640 with 10% FBS media to each well of the xCELLigence E-96 plate. MCF7 or B80-T5 were seeded at a density of 20,000 or 10,000 cells/well in a volume of 100 µL RPMI1640 with 10% FBS media and allowed to passively adhere on the electrode surface. IFNγ was added to the media to achieve the final concentration of 10 ng/mL. After 6 hours, 10 µM SGC3027 (Sigma Aldrich) was added, or 0.1% DMSO as a negative control. Eighteen hours after adding SGC3027, the culture medium was refreshed (removing IFNγ, SGC3027 and DMSO) and T cells were added at an E:T ratio of 1:1. Cytolytic rates for samples exposed to T cells were normalised using the cell index values from unexposed controls.

## Results

### Selection of genes for the targeted library

To build the custom CRISPR libraries, we selected the target genes at 205 fine-mapped breast cancer risk GWAS signals predicted using the INQUISIT pipeline^3^, genes from Transcriptome-Wide Association Studies (TWAS) and expression quantitative trait loci (eQTL) studies of breast cancer risk ^4^ (Figure 1a, Table S1). We designed five sgRNAs for each gene, and included 1000 negative control sgRNAs targeting the *AAVS1* region, 1000 sgRNAs targeting 193 core essential genes, and 27 positive control genes known to confer resistance to T cell–mediated killing (Table S2).

**Figure 1.**
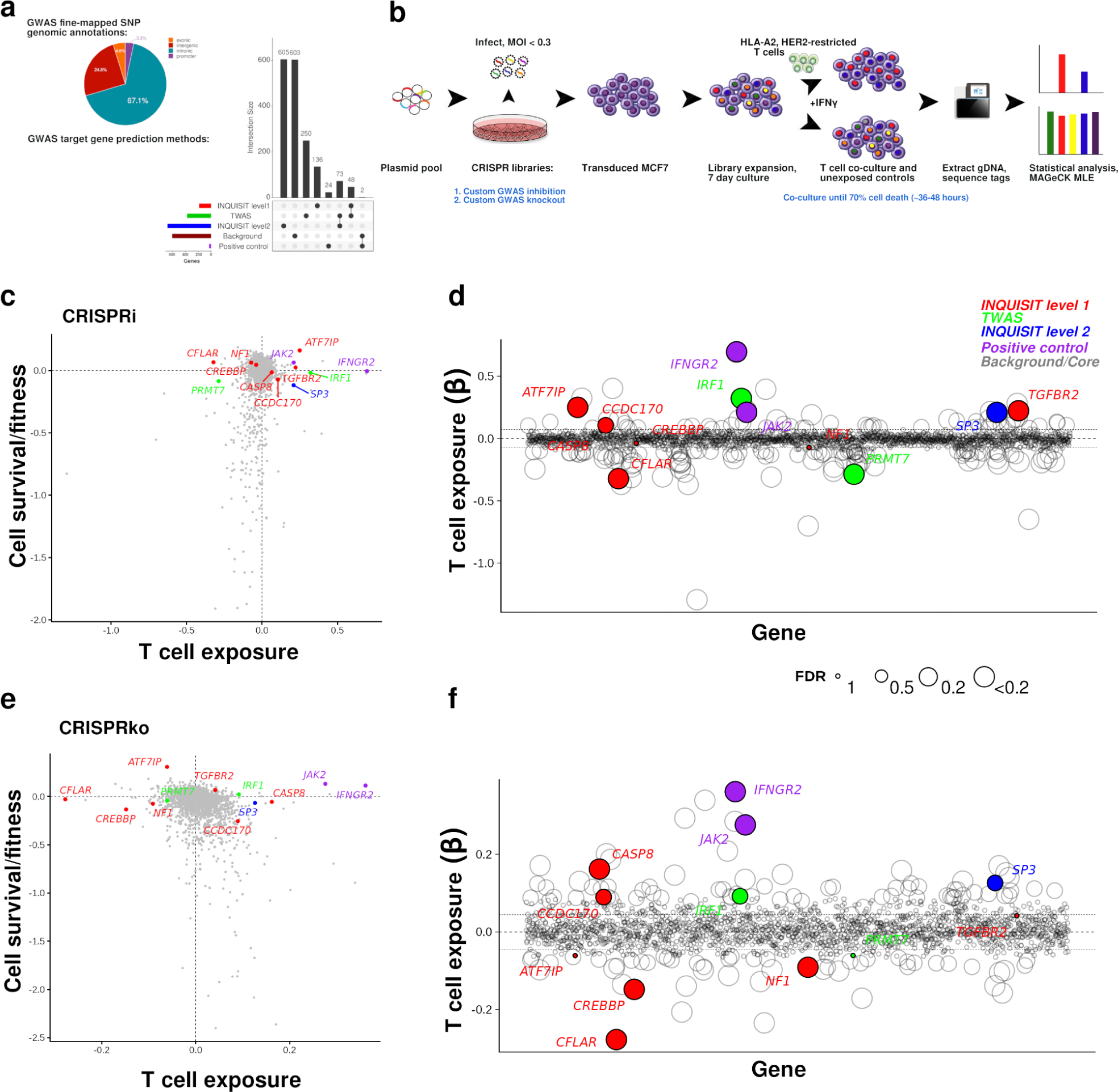
Gene selection, experimental design and workflow to identify candidate breast cancer risk genes which affect cytotoxic T cell-mediated MCF7 tumor cell killing. Genes from breast cancer GWAS were selected from previous studies (**a**). MCF7 cells were infected with custom GWAS knockout and inhibition libraries before co-culture with T cells. Genomic DNA was extracted from surviving cells and the difference in guide abundance in the unexposed and co-cultured groups analysed with MAGeCK (**b**). Results comparing the β effect of T cell exposure to the β effects attributed to cell fitness in CRISPR inhibition (**c**) and knockout (**e**) screens. Negative β values are associated with genes that are essential for cell survival and/or T cell evasion. Labelled genes were selected for validation. The circle sizes in (**d**) and (**f**) indicate adjusted significance levels and lines show ±1 SD. Label colours indicate the gene prediction method (high confidence INQUISIT red, moderate confidence INQUISIT blue, TWAS green, positive controls purple).

### Optimization of the CTL assay

In order to identify the best cell line for the screen, we compared the cytolysis efficacy of HER2-restricted T cells against four HLA-A2 positive breast cell lines, MCF7, SKBR3, MDA-MB-231 and B80-T5 (Figure S1a). We noted that although only SKBR3 has copy number amplification and very high expression of HER2, all other cell lines express moderate levels of cell surface HER2 protein (Figure S1b). While MCF7 are not classified as HER2 amplified cells, they were most sensitive to HER2-restricted T cells with a 20 hour delay in onset, followed by an almost linear killing curve, achieving 90% lysis after a 52 hour co-culture period. In comparison, SKBR3 and MDA-MB-231 were more resistant to HER2-restricted T cells with the cytolysis rate plateauing at about 30% (Figure S1a).

Consistent with previous studies ^23^, we found that MCF7 treated with 10 ng/ml of IFNγ for 24 hours robustly up-regulated HLA-ABC expression (Figure S1c). Such increased HLA class I expression is associated with increased efficiency of antigen processing and presentation ^21,28,33–36^. We also compared T cell killing efficiency by altering E:T ratios from 0.25 to 8, and showed that a ratio of 1:1 exerted intermediate immune pressure (Figure S1d), suitable for the phenotypic screen endpoint. By comparing T cell killing efficiency of HER2-restricted T cells versus T blasts, we showed the specificity of our assay (Figure S1e). Following this optimization, we used the following co-culture conditions for the screen: HER2-restricted T cells and target MCF7 cells at a 1:1 ratio, with target cell IFNγ pretreatment, and an endpoint of 70% cytolysis in the parallel xCELLigence assay.

### Identification of candidate breast cancer risk genes which regulate sensitivity to T cell cytotoxicity

To gain a comprehensive view of breast cancer GWAS targets associated with immune evasion, we performed functional CRISPR screens, using custom CRISPRi and CRISPRko libraries (Figure 1b). Following independent library infections in triplicate cultures, cells were grown for seven days to deplete cells with essential gene perturbations. MCF7 cells were co-cultured with HER2-restricted T cells and when 70% cytolysis was observed the cells were harvested and genomic DNA used for quantification of sgRNA abundance. We compared sgRNA abundance between the co-cultured cells and unexposed controls using the MAGeCK pipeline ^37^. Sequencing analysis revealed sgRNA library mapping rates >90% and Gini index <0.1 for all sample libraries. Consistent read distributions indicated that guides were adequately represented at all stages of the workflow. We further assessed screen quality by examining the levels of sgRNAs targeting 193 core essential genes. Compared to early experimental timepoints (day 3), the log read count of essential genes relative to non-essential genes was substantially left-shifted in later time points in inhibition and knockout screens (Supplementary Figure 2a and c). We further distinguished between candidate CTL evasion effects and genes with survival promoting functions in MCF7. We retrieved fitness genes from the Cancer Dependency Map Project and designated MCF7 essential genes as *Chronos score* < -1 (892 genes, of which 226 were in the screen library). We found that mean sgRNA abundances for the genes known to contribute to MCF7 survival in culture were again left-shifted relative to non-essential screen genes (Supplementary Feig 2b and d).

We performed CRISPRi and CRISPRko screens with custom libraries and analysed sgRNA abundances with MAGeCK (Figure 1c-f). We used the maximum likelihood estimation (MLE) approach to model experimental conditions and compute an effect score (‘β’) which reflects the extent of selection (Table S6). To account for fitness effects, we modelled a variable to account for general effects on cellular fitness in culture in our screen data. We set an adjusted significance threshold of 0.2 to identify candidate breast cancer risk genes that act as ‘sensitizers’ to T cell killing (β > 0, relative sgRNA enrichment in surviving cells), and ‘resistors’ that enable MCF7 cells to evade T cell cytotoxicity (β < 0, sgRNA depletion). As expected, negative controls had no effect on cytotoxicity. We observed positive correlation between screens (ρ = 0.25; *P* < 2.22×10^−16^), indicating that gene perturbation using different modalities resulted in concordant effects (Figure S3). We identified 88 genes which were significantly enriched or depleted upon T cell-mediated tumor cell killing (FDR < 0.2, Table S7). Positive control genes were detected in the inhibition (*CD58, IFNGR1, IFNGR2, JAK1, JAK2, NLRC5* and *TAP1*) and knockout screens (*CD58, HLA-A, IFNGR1, IFNGR2, JAK1, JAK2*, and *TAP1*), and were associated with positive β values (Table S6) indicating that perturbation induced resistance to immune killing. Eleven genes were detected in both screens.

To differentiate between genes essential for cell survival and probable CTL genes, we examined sgRNA abundances across all treatment conditions compared to baseline. We found 9 and 26 genes (FDR < 0.05) with negative β values for the fitness variable (indicating depletion over time in culture) in the knockout and inhibition screens, respectively, which are likely to be genes depleted due to effects on cell survival rather than CTL evasion. These genes include GWAS targets *CCND1, ESR1, GATA3* and *EWSR1*. As expected, essential genes were enriched in core pathways such as DNA repair and proteasome degradation pathways. Using data from the DepMap Project, 31 genes known to affect MCF7 fitness were not considered as putative CTL evasion genes. The final filtered list of candidate GWAS target genes with an effect on T cell killing comprised 20 resistors and 13 sensitisers. All 33 genes were predicted to be target genes at GWAS signals (25 by INQUISIT and 8 by TWAS). Using the library content as background, analysis using the Enchichr tool showed the 33 CTL genes were enriched for pathway terms including Apoptosis, IL6 signalling and Interferon response.

### Validation of candidate CTL evasion genes

To validate a subset of these 33 candidate breast cancer CTL genes identified in these screens, we performed single gene perturbation experiments with CRISPRko and CRISPRi. The selection criteria for this subset was genes were 1) predicted with high confidence by INQUISIT (*CFLAR* and *CREBBP*) or from TWAS (*IRF1* and *PRMT7*); 2) predicted with high confidence, and previously shown to be regulated by breast cancer risk variants (*ATF7IP, CASP8*, and *CCDC170*). In addition to these seven genes, we included two positive control genes (*IFNGR2* and *JAK2*) in validation assays.

To perform the single gene validation, we infected MCF7 with the top two scoring CRISPRko or CRISPRi sgRNAs from the custom libraries (except for *CASP8* which did not score in the CRISPRi screen, and so we re-designed the CRISPRi sgRNAs), followed by puromycin selection for 7 days. Gene perturbation was assessed by Western blotting (Figure S4), or quantitative real-time PCR (Figure S5). On day 7, the lines were pretreated for 24 hr with 10 ng/ml IFNγ and co-cultured with T cells at an E:T ratio of 1:1 while their growth was monitored by xCELLigence. The rate of cytolysis was monitored until 70% MCF7 lysis was achieved upon exposure to HER2-restricted T cells. The relative cytolysis in the xCELLigence assay was determined relative to cells infected with a non-targeting control guide (sgRFP).

As expected, inhibition or knockout of the positive control genes, *IFNGR2* and *JAK2*, rendered MCF7 cells more resistant to the HER2-restricted T cells (Figure S6). We validated the role of *ATF7IP, CASP8, CCDC170*, and *IRF1* as sensitizers, and of *CFLAR, CREBBP* and *PRMT7* as resistors. CRISPRi did not knockdown *CREBBP* expression (Figure S2), but the CRISPRko results validated it as a resistor.

### Identification of IRF1 and CASP8 regulatory variants

*IRF1* was included in our CRISPR library on the basis of TWAS analysis ^38^ using data from multiple eQTL studies. However, the direction of expression associated with breast cancer risk is not consistent between sources. We therefore looked for candidate risk variants that might explain the association. The GWAS catalog contains a variant ∼7 kb upstream of the *IRF1* promoter (rs736801, *P* = 8×10^−8^) in a putative regulatory element annotated as and Active Enhancer in MCF7 cells in the RegulomeDB database ^39^ and a distal enhancer-like signature ENCODE SCREEN based on H3K4me1 and H3K27ac ChIP-seq signals (Figure 2a). To determine whether this region controlled *IRF1* expression, we performed CRISPRi in MCF7 cells. Using qPCR, we found a 2-fold reduction in *IRF1* expression suggesting direct distal regulation by this upstream putative regulatory element (PRE). We hypothesised that reduced expression mediated by perturbation of this PRE would mimic direct *IRF1* gene knockout and lead to reduced T cell-mediated MCF7 killing. We performed CTL assays using MCF7 cells with *IRF1* gene deletion and PRE modulation. Compared to T cell mediated cytolysis of control cells, we observed significantly reduced killing of PRE-modulated target cells (Figure 2b). These results show that *IRF1* gene expression can be inhibited via a distal regulatory element to levels which influence immune interactions.

**Figure 2.**
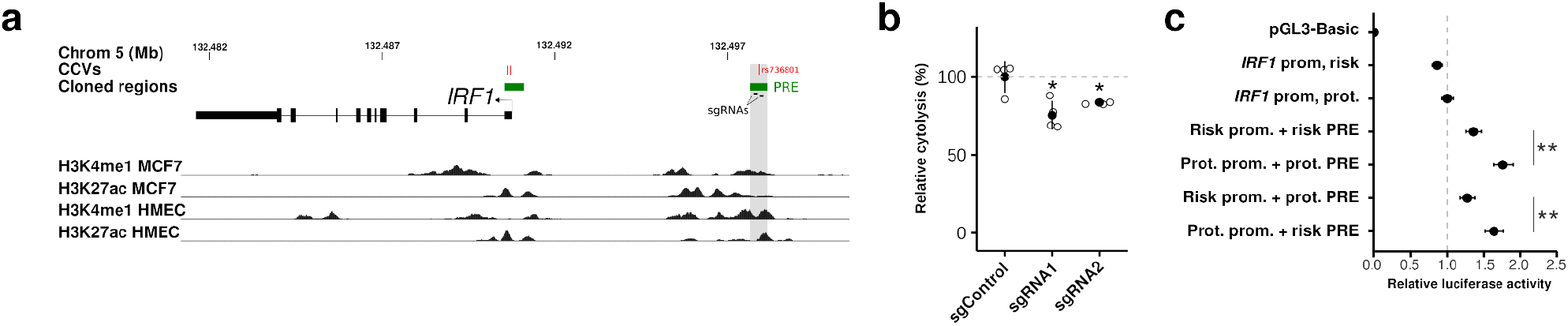
Regulation of *IRF1* by an upstream non-coding variant. a) shows the genomic landscape around *IRF1* with positions of the candidate risk variants (red), cloned regulatory regions (green) and histone ChIP-seq data for MCF7 and HMEC H3K4me1 and H3K27ac marks from ENCODE. b) CRISPRi of the region around rs736801 using sgRNAs as depicted in (a) reduced cytolysis in the CTL assay. Filled circles denote the mean and whiskers represent standard deviations. Significance was computed with two-sample T tests, using Benjamini-Hochberg correction for multiple testing. All experiments comprised n > 3. (\**P* < 0.05). c) Reporter constructs containing promoter and putative enhancer regions corresponding to green regions in (a) were transfected into MCF7 cells, revealing the protective haplotype was associated with increased *IRF1* promoter activity. Filled circles denote sample means and whiskers represent 95% confidence intervals. Significance was computed using two-way ANOVA followed by Dunnett’s test for multiple comparisons. All experiments comprised n ≥ 3. (\*\**P* < 0.01).

We next tested allele-dependent effects of the candidate causal variant (CCV) rs736801 on *IRF1* promoter activity. We cloned the *IRF1* promoter along with PREs containing reference and alternate (risk) alleles into luciferase reporters. Two correlated SNPs (rs11242115 and rs10900809, r^2^ = 1 with rs736801 in 1000 Genomes CEU population) overlapping *IRF1* exon 1 were included in the promoter fragments. We found that the PRE drove a ∼2-fold increase in promoter activity (Figure 2c). The risk haplotype constructs(rs736801-T, rs11242115-C, rs10900809-A) lead to significantly diminished promoter activity compared to the protective haplotype (*P* = 0.002; Figure 2c). This effect is consistent with the risk allele of rs736801 altering PRE activity and reducing *IRF1* expression. These molecular analyses support a consistent direction between genetic risk and the effect of lower IRF1 levels on the T cell evasion phenotype.

We also used reporter assays to evaluate rs3769821, a fine-mapped CCV in the promoter of an isoform of *CASP8*. This CCV lies in an intronic region characterised by regulatory signatures (H3K4me1 and H3K27ac) in MCF7 and HMEC cells (Figure 3a and b). We generated clones with risk and protective alleles within the *CASP8* promoter to drive expression of the luciferase reporter ^40^. We transfected MCF7 cells and detected a 30% reduction in luciferase activity with the risk allele (*P* = 0.019; Figure 3c). This effect suggests that the breast cancer risk allele of rs3769821 leads to reduced *CASP8* expression, an effect consistent with lower CASP8 levels being associated with diminished T cell killing of target tumor cells.

**Figure 3.**
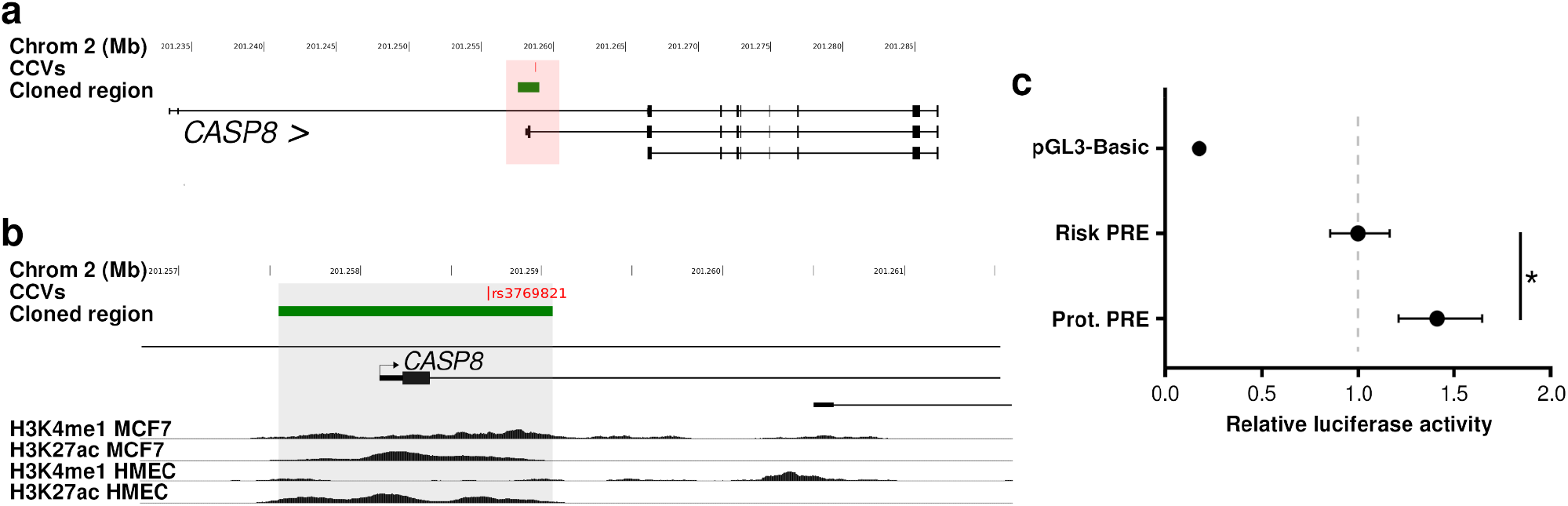
Regulation of *CASP8* by a promoter variant. (a) Genomic landscape and zoomed in view (b) around the promoter of an alternate *CASP8* promoter. Reporter constructs containing the *CASP8* promoter with risk or protective alleles of rs3769821 were transfected into MCF7 cells (c), where filled circles denote sample means and whiskers represent 95% confidence intervals. Significance was computed using two-way ANOVA followed by Dunnett’s test for multiple comparisons. All experiments comprised n ≥ 3. (\**P* < 0.05, \*\**P* < 0.01).

### Immune signatures in clinical samples and patient outcomes

*PRMT7* deletion is associated with increased CTL activity *in vitro*, so we examined associations between gene expression and breast cancer patient outcomes. In Luminal A cases in the TCGA cohort, we observed significantly reduced progression-free interval (PFI) in cases with elevated *PRMT7* expression (*P*_log-rank_ = 0.043, Figure 4a), an effect which was validated in Luminal A tumors in the Metabric cohort ^30^ (*P*_log-rank_ = 0.035, Figure 4b). A concordant trend was seen for overall survival in both cohorts (Supp Fig 7). We hypothesised that patient outcomes may be dependent on the interaction between *PRMT7* expression and levels of infiltrating CD8 cells. Stratifying TCGA Luminal A tumors by CD8+ T cell infiltration levels revealed a reduction in PFI survival time in patients with high *PRMT7* expression and low levels of CD8+ compared with low PRMT7 and high CD8+ levels (*P* = 0.0058; Figure 4c). These findings suggest that high *PRMT7* levels in tumors is associated with reduced patient survival through an interaction with CD8+ T cell activity. This effect is consistent with the resistor activity observed in our T cell killing assays and indicates that PRMT7 inhibition may be clinically beneficial.

**Figure 4.**
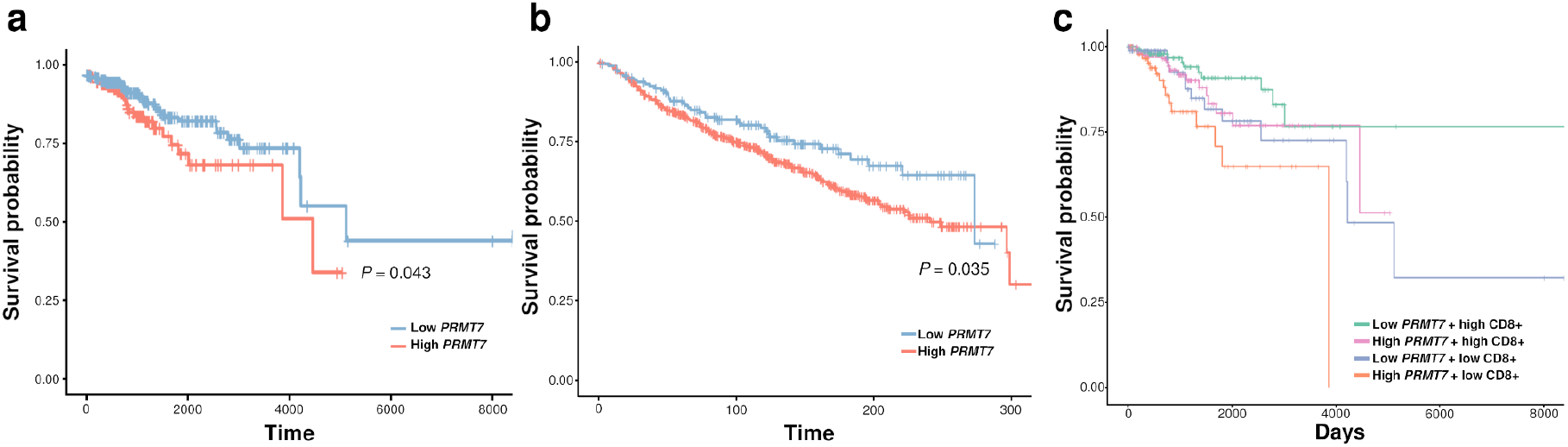
*PRMT7* expression is associated with survival in patients with Luminal A breast cancer. High *PRMT7* expression is significantly associated with reduced progression-free interval (PFI) in the TCGA cohort (**a**) and relapse-free survival (RFS) in the Metabric cohorts (**b**). *PRMT7* expression is associated with reduced PFI in TCGA Luminal A tumors with low CD8+ T cell infiltration compared with tumors with high CD8+ T cell levels (**c**). *P* values were computed using log-rank tests.

We next examined the relationship between expression of *PRMT7* and immune signatures in breast tumor samples using data from The Cancer Genome Atlas ^31^. *PRMT7* levels were negatively correlated with CD8+ levels (Figure 5a. Furthermore, this association was restricted to Luminal A and B tumors (ρ = -0.262, *P* = 1.49x10^−9^ and ρ = -0.16, *P* = 0.023, respectively). We noted a lack of correlation with tumor purity (*P* = 0.446, Figure 5b), indicating that *PRMT7* expression was not specific to tumour or stromal cells.

**Figure 5.**
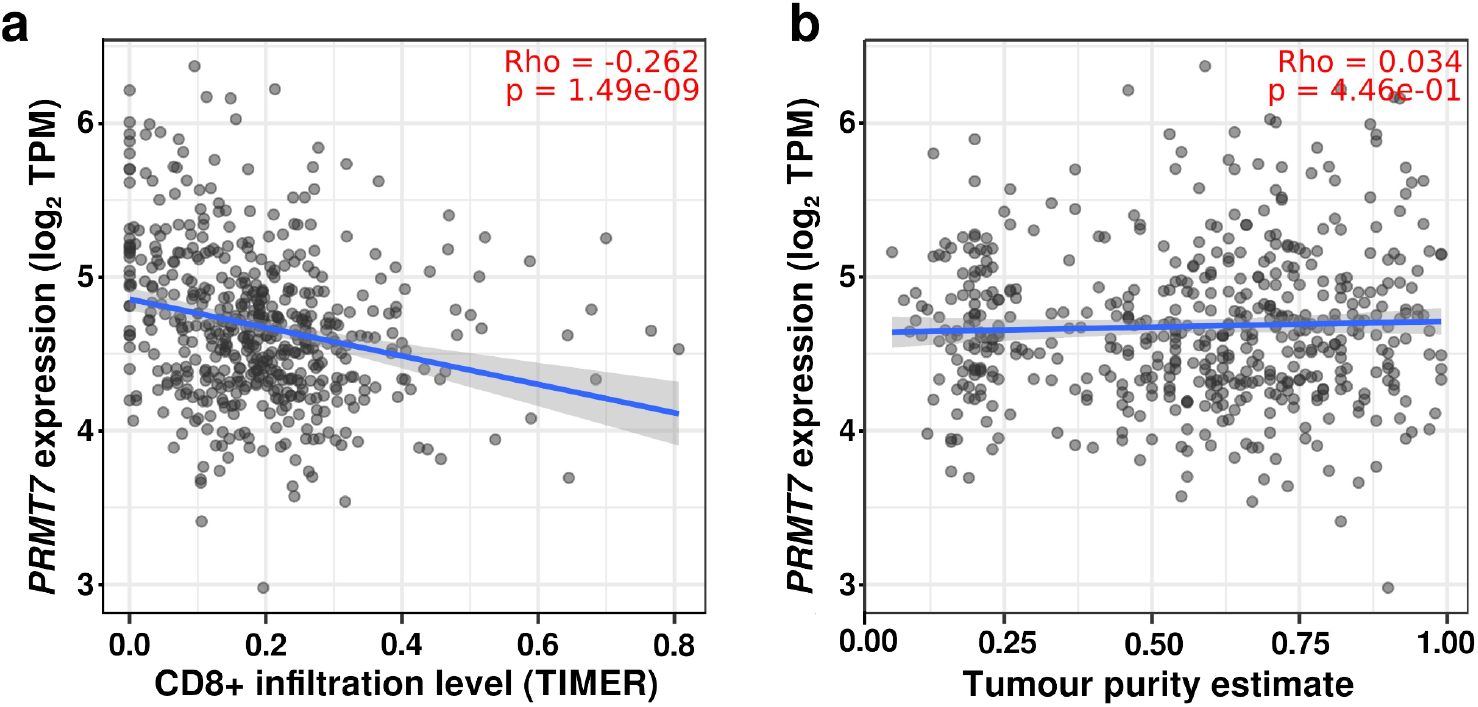
Gene expression and levels of CD8+ T cell infiltration in TCGA breast tumors. *PRMT7* expression is inversely correlated with CD8+ T cell levels in Luminal A tumors (**a**), and is not correlated with tumor purity (**b**).

### PRMT7 pharmacological inhibition in vitro

In light of the association between lower *PRMT7* levels and improved patient survival, reducing PRMT7 activity may represent a novel breast cancer treatment strategy. Importantly, inhibition of PRMT7 with a selective small molecule inhibitor, SGC3027, can sensitize melanoma to immune checkpoint blockade ^41^. In addition to PRMT7 perturbation in MCF7 cells, we repeated PRMT7 knockout and inhibition in the immortalized, normal breast cell line, B80T5. We achieved >90% reduction in PRMT7 protein levels (Figure 6b), and observed up to a 2-fold increased rate of T cell-mediated lysis in B80T5 target cells with both modes of perturbation (Figure 6c and d). We next investigated the effect of PRMT7 pharmacological inhibition with SGC3027. Following pre-treatment of target MCF7 or B80T5 (18 hours), inhibition of PRMT7 significantly increased the rate of T cell-mediated cytolysis (Figure 6e and f). These results demonstrate that genetic and pharmacological inhibition of PRMT7 enhances the degree of immune killing in multiple breast cell contexts.

**Figure 6.**
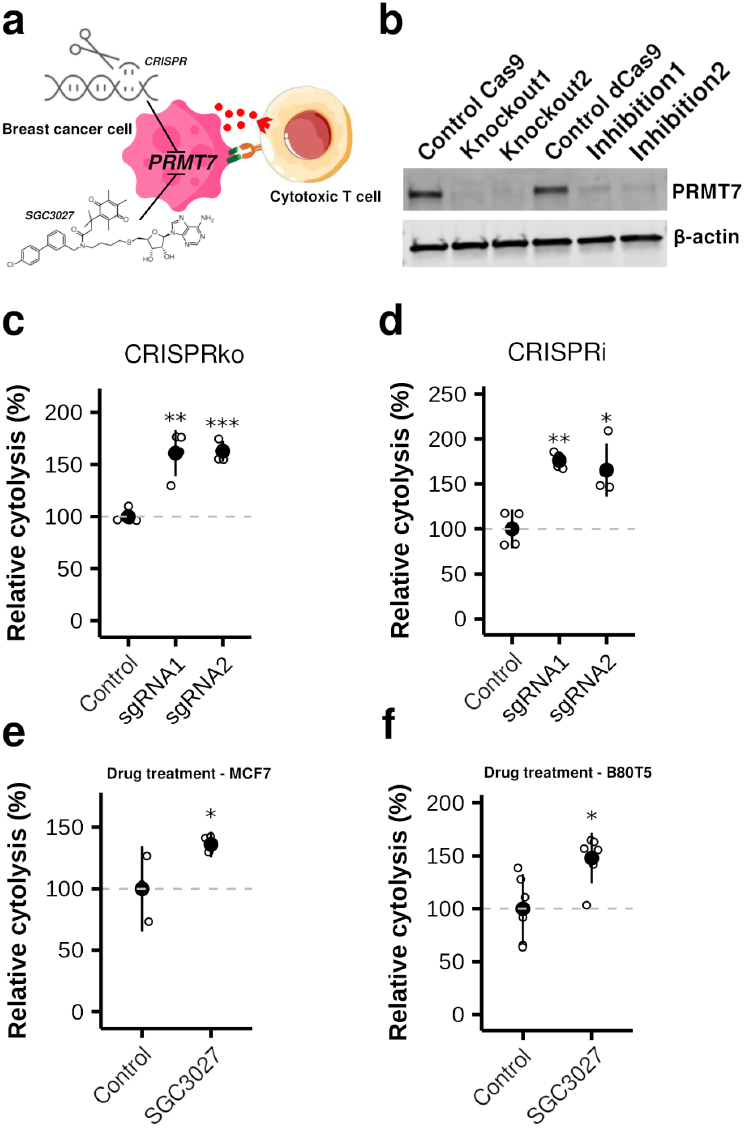
Inhibition of PRMT7 activity. Genetic perturbation and pharmacological inhibition of *PRMT7* alters response to T cell killing (**a**). Depleted PRMT7 protein levels in independent B80-T5 lines up CRISPRko and CRISPRi (**b**). Relative cytolysis (compared to cells with non-targeting guide) in B80-T5 following CRISPRko (**c**) or CRISPRi (**d**). PRMT7 inhibition with SGC3027 enhanced T cell-mediated killing of both MCF7 (**e**) and B80T5 (**f**). Filled circles denote sample means and whiskers represent standard deviations. Significance was computed with two-sample T tests, using Benjamini-Hochberg correction for multiple testing. All experiments comprised n > 3. (\**P* < 0.05, \*\**P* < 0.01, \*\*\**P* < 0.001, \*\*\*\**P* < 0.0001).

## Discussion

Genome-wide association studies have been very successful at identifying loci associated with breast cancer risk, with over 220 identified so far ^1,2^. Most of the candidate causal variants identified through fine-mapping lie in non-coding regions, and most likely act by regulating the expression of one or more nearby genes ^3^. One of the major goals of GWAS is to find target genes and proteins that might lead to new strategies for prevention or treatment. This is particularly important since several studies have shown that the likelihood of a drug getting to market is approximately doubled if it has genetic support ^42,43^. However, identifying the target gene(s) at a locus, which has been done for less than 20 loci by chromatin conformation analysis and luciferase assays ^44–47^, is time consuming and does not address the issue of molecular pleiotropy: it is important to know which target gene(s) can impact on a phenotype relevant to the disease or trait of interest. To complement studies of individual risk loci, pooled functional screens for follow up of GWAS findings for coronary artery disease, lung disease, blood traits, type 2 diabetes and height ^48–52^. We have undertaken functional CRISPR screens to enable the identification of genes near GWAS signals that impact the hallmarks of cancer.

Of the 33 genes we identified in our CTL screens, 18 had not previously been found through a re-analysis of tumor immunity-associated functional screens ^53^. Consistent with our findings, *CASP8* has been reported as a sensitizer previously, and *CFLAR* and *CREBBP* as resistors ^23^. In contrast, *ATF7IP* has been previously shown to be a resistor in a murine melanoma model ^54^, and *IRF1* and *PRMT7* have been variably found to be resistors or sensitizers ^23,41^. Lawson et al ^23^ carried out CTL screens in two murine triple negative mammary carcinoma cell lines and identified *IRF1, CFLAR, CREBBP, ATF7IP, CASP8* and *PRMT7* as hits, although not always in the same direction as we found in our screens. Notably, the murine cell lines used were both of the “Triple Negative” histological subtype, which suggests that cell type specific differences may influence immune phenotypes.

*PRMT7* was included in our screen because it was found through a transcriptome-wide association study of breast cancer risk ^55^, where risk was associated with increased predicted expression. This oncogenic effect is consistent with an inverse relationship between expression and survival in TCGA and Metabric cohorts. PRMT7 has been shown to promote breast cancer metastasis ^56^, but a recent report showed that its inhibition with the small molecule inhibitor, SGC3027, ^41^ stimulates anti-tumor immunity and sensitizes melanoma to immune checkpoint blockade. Here, we show that in the context of reduced breast cancer patient survival, high *PRMT7* expression levels are associated with lower levels of CD8+ infiltration. Our results showing that SGC3027 increases T cell-mediated cytolysis of MCF7 and B80-T5 breast cells *in vitro*, suggest that it may also be effective to control breast tumorigenicity *in vivo*, with or without immune checkpoint inhibitors.

A major challenge for post-GWAS analyses is to link DNA variation to target gene function. We found such evidence for several CTL genes. *IRF1* (Interferon Regulatory Factor 1) was included in our screen because it was found through a transcriptome-wide association study of breast cancer risk ^38^. Subsequent investigation revealed a variant at this locus, rs736801, was associated with risk at sub genome-wide significance levels (*P* = 8 x 10^−8^) in large breast cancer GWAS ^2^. Our molecular analyses suggest this risk allele directly reduces *IRF1* expression. IRF1 is a critical mediator of the IFNγ signalling transcription factor family ^57^ which have diverse roles in the gene-regulatory networks of the immune system. IRF1 controls expression of antigen presentation machinery responsible for loading antigens on MHC class I molecules ^58^ thereby facilitating T cell cytotoxicity. However, others have found that IRF1 inhibits antitumor immunity through the upregulation of PD-L1 in tumor cells ^59^, which would be consistent with the resistor role found in other functional screens.

*CASP8* and *CFLAR* are both high confidence prediction target genes ^3^ at the same breast cancer risk region at the chromosome 2q33.1 locus. CASP8 is most commonly recognized for its role in apoptosis but it also regulated B and T lymphocyte activation, and macrophage differentiation and polarisation^60^. The role of CFLAR (CASP8 And FADD Like Apoptosis Regulator) in resisting T cell cytotoxicity that we and others have found through functional screens may be because high tumoral expression can inhibit cell death induced by immune effector cells via the FAS and TRAIL death receptors ^61^. Indeed, FAS Ligand (FASLG) and TRAIL expressed by cytotoxic T lymphocytes and natural killer cells may exert immunological selection pressure for tumors with high *CFLAR* expression. In support of this, overexpression was sufficient to permit establishment of tumors in immune-competent mice by inhibiting Fas-dependent cell death ^62^. Interestingly, *CASP8* and *CFLAR* were detected as a sensitizer and a resistor in our screens, consistent with their known opposing functions. Further molecular work is needed to understand the relationship between breast cancer risk-associated genetic variation and these genes. Previous studies have shown genetic regulation of CASP8 expression, but a complex array of GWAS signals cover this locus. An outstanding question remains as to whether independent genetic signals control *CASP8* and *CFLAR* expression in opposing directions.

In conclusion, we have identified a set of breast cancer risk genes which function in cancer cells by regulating their sensitivity to the cytotoxic T lymphocytes. Follow up of these findings, particularly PRMT7 for which a small molecule inhibitor is available, might lead to novel therapies for treatment and prevention. Our findings highlight the value of pooled functional CRISPR screening for identifying genes that operate through hallmarks of cancer which are less studied in the context of breast cancer risk. This approach may help to expedite the translation of GWAS to novel treatments and risk reduction strategies.

## Acknowledgments

This work was funded by the National Health and Medical Research Council Ideas Grant 1185615 and Investigator Grant 2020/GNT1194914, and by an award for research excellence from GSK to GCT. We thank Pauline Crooks for performing venipuncture. Metabric data was downloaded from https://ega-archive.org/dacs/EGAC00001000484 with permission from the data access committee.

## Author contributions

W.S., Y.L., D.B., A.C., M.M., N.T., conducted the experiments; J.Bu., C.X., Z.P., S.T., and C.S. provided advice; J. Be., M.R. and L.G-J. performed the statistical analyses; W.S., J.R., J.Be. and G.CT.. designed the experiments and wrote the paper

